# Tet Transgene Activation is Disrupted in Lipogenic Triple Negative Breast Cancer Cells

**DOI:** 10.1101/2024.10.18.619156

**Authors:** Ashley Townsel, Yifei Wu, Maya Jaffe, Cara Shields, Karmella A. Haynes

## Abstract

An important challenge for mammalian cell engineering is the unexpected response of transgenes to native transcriptional regulation pathways. One transgene can show different levels of expression at different genomic sites, in different cell types, and under different growth conditions. Collisions between transcription and DNA replication, heterochromatin encroachment, and viral defense have been linked to transgene silencing. In this study we identify fatty acid metabolism as another mediator of transgene behavior. Adipocyte secretome-induced lipogenesis in epithelial breast cancer cells was accompanied by the loss of expression from a Tet-TA regulated *pCMV*-AmCyan fluorescent protein (CFP) transgene. Transcription profiling verified the activation of lipid droplet biosynthesis genes, and revealed repression of loci at every chromosome, consistent with the idea that lipogenesis affects the availability of substrates and cofactors for global chromatin remodeling. Pre-induction of *pCMV* prevented full transgene silencing during lipogenesis. Our results provide new insights into the influence of the lipogenic epigenetic state on transgene behavior.

## INTRODUCTION

Transposon systems are indispensable for cell engineering, as they allow the stable integration of transgenes into the host cell genome, ensuring heritable long-term expression in proliferating cell populations. The expression of transgenes can differ between cell types and can be affected by both intrinsic and extrinsic factors, making transgene expression unpredictable. Competition between the transcriptional machinery needed to express transgenes can interfere with endogenous gene expression patterns and cause silencing ^1^. For example, native genes with methylated CpG islands can recruit histone deacetylases to nearby transgene insertion sites and compete with mechanisms of transgene expression that rely on histone acetylation ^2,3^. Furthermore, the activity of histone methyltransferases at heterochromatin can spread to nearby transgene insertion sites and cause gene silencing ^4,5^. These mechanisms of repression force transgene expression to default to native chromatin regulation pathways. Transgene silencing can also be a response to the transgene itself. The cellular recognition of viral components in transposon systems can cause changes in chromatin states that initiate repressive defense systems against transgene elements ^6–8^.

Chromatin states can also change in response to environmental stimuli and culture conditions, which alter levels of metabolites involved in cellular metabolism and respiration. One type of stimulus known to induce substantial changes in cellular nutrient storage and consumption is adipocyte signaling. Adipocyte secreted factors including insulin, CCL5, and apelin interact with receptors that stimulate mTOR signaling in neighboring cells. One of the major transcription factors that acts downstream of this signaling pathway is SREBP1, which translocates into the nucleus and activates the expression of metabolic enzymes and other proteins involved in fatty acid synthesis (lipogenesis), transport, and storage ^9^. Their associated enzymatic activities increase or decrease pools of metabolites that serve as substrates and cofactors for chromatin remodeling. Major epigenetic metabolites that are linked to fatty acid processing include acetyl-CoA generated from fatty acid beta oxidation, alpha-ketoglutarate (α-KG) and flavin adenine dinucleotide (FAD+) which are generated or consumed (respectively) the TCA cycle, and nicotinamide adenine dinucleotide (NAD+) generated by the salvage pathway ^10–13^. Acetyl-CoA is the sole substrate for chromatin remodeling by nuclear histone acetyltransferases (HATs), which supports active gene transcription. A close relationship between global levels of histone acetylation and acetyl-CoA production has been demonstrated in various human cancer cell types, other mammalian cells including adipocytes, endothelial cells, mesangial cells, and immune cells ^14^. In yeast, inhibition of ACC1 results in increased global histone acetylation ^15^. Therefore a decrease in the availability of acetyl-CoA might lead to a global loss of acetylation, and epigenetic repression of endogenous genes and transgenes.

In this study, we observed changes in the expression of a chromosomally integrated sleeping beauty based transgene, pSBtetTA, as breast cancer epithelial cells became lipogenic in response to adipocyte-secreted factors (ASFs) *in vitro*. The general-purpose multi-gene expression system pSBtetTA allowed us to observe epigenetic effects at two different promoters, one doxycycline-inducible promoter (*pCMV*) regulated by a VP16 fusion regulator (Tet-TA), and a separate constitutive promoter (*pRPL13a*) ^16^. RNA-seq profiling verified the activation of fatty acid metabolism and storage genes in lipogenic cells, and revealed epigenetic silencing of other genes throughout the genome. We used imaging and flow cytometry to determine that Tet-TA regulated expression is strongly repressed in lipogenic breast cancer cells but not in non-lipogenic breast cancer or HEK293 cells. To determine how transcription initiation and active transcription might be distinctly affected by the lipogenic epigenetic state, we induced lipogenesis before, during, and after transcriptional activation with doxycycline. We observed that inducing the lipogenic epigenetic state prior to transcriptional activation blocked transgene expression more effectively than when lipogenesis was induced after transcriptional initiation.

## MATERIALS AND METHODS

### Cell Culture

Standard media for each cell line was as follows: BT-549 cells (ATCC #HTB122) - RPMI-1640 (ATCC #30-2001) with 10% tetracycline system approved fetal bovine serum (tsFBS, Thermo #A4736401), and 0.8 ug/mL insulin (Thermo #12585014); HEK293 cells (ATCC #CRL-1573) - DMEM High Glucose (Thermo #11965092) with 10% tsFBS; OP9 cells (ATCC #CRL-2749) - alpha-MEM (Corning #15-012-CV) with 20% fetal bovine serum (FBS, Bio-Techne #S11150), and 1% penicillin-streptomycin (Thermo Fisher #15140122); Medium for stromal OP9 cell propagation - DMEM (Hyclone #SH30243.01) with 10% FBS and 1% penicillin-streptomycin; Insulin Oleate Medium (IOM) for OP9 differentiation - alpha-MEM with 1.52 mM sodium oleate (Sigma #O7501) in methanol, 1.8% fatty acid free bovine serum albumin (Sigma #A6003) in 1X PBS, 0.2% FBS (Bio-Techne #S11150), 175 nM insulin (Sigma #I6634), 1% penicillin-streptomycin. IOM was prepared as described in **Supplemental Methods**. Cells were grown at 37°C, with 5% CO_2_ in a humidified incubator unless otherwise noted. 1X DPBS without calcium and magnesium (Corning #20-031-CV) and 0.25% trypsin-EDTA (Thermo #25200056) was used for cell washing and harvesting. For long term storage, 1 - 2×10^6^ cells were frozen in 1 mL 10% DMSO in tsFBS or FBS at -80°C for 2 days, then transferred to -150°C (liquid nitrogen, LN2).

### Preparation of Adipocyte-Conditioned Media (ACM)

OP9 cells were seeded at a density of 5×10^5^ and cultured in OP9 stromal propagation medium for 24 hours. The medium was replaced with 10 mL IOM, cells were cultured for 72 hours, and the supernatant adipocyte-conditioned media (ACM) was collected and stored frozen -20°C) as 6 mL aliquots. ACM was diluted to 50% with unconditioned medium (UCM) and used for the Oil Red O staining and RNA-seq experiments, while 75% ACM/UCM was used for all other experiments.

### Oil Red O Staining

Oil Red O stock solution was prepared as 0.35% (g/mL) Oil Red O (Sigma-Aldrich #O0625) in 100% isopropyl alcohol and passed through a 0.22 μm filter. Oil Red O working solution was prepared with 3 parts Oil Red O stock solution and 2 parts distilled water, passed through a 0.22 μm filter and incubated at room temperature for 10 minutes. TNBC cells were seeded at 5×10^4^ cells/mL in a 6-well plate and grown overnight before replacing 2 mL culture medium with either UCM or 50% ACM for 48 hours. The cells were washed with 1X DPBS, fixed with 4% paraformaldehyde (Thermo Fisher #J61899-AK) for 30 minutes, rinsed with distilled water, and equilibrated with 60% isopropanol for 2 minutes at room temperature. Cells were stained with the Oil Red O working solution and incubated at 37°C for 10 minutes. To visualize nuclei in non-transgenic cells, samples were counterstained with 1 mL Hematoxylin (Vector Laboratories #H-3404-100) per well for 30 seconds, and rinsed multiple times with deionized water. All samples were covered with 1X DPBS and imaged at 200x.

### RNA-Seq

BT-549 cells were seeded at 1×10^6^ cells/mL in a 6-well plate and grown in UCM or 50% ACM, with 3 replicates per condition, for 24 hours. Media was removed, adherent cells were lysed directly in the wells, and RNA was extracted and purified as instructed by the RNeasy Mini kit (Qiagen #74104). Frozen RNA (-80°C) was submitted to Novogene for library preparation (polyA-enriched) and next generation sequencing with the following parameters: paired-end, 150 bp, Q30 ≥85%. Raw reads were trimmed with Trim Galore and aligned via seed searching using STAR ^17^ with the most recent human genome (GRCh38/hg38). mRNA levels were calculated with RSEM ^18^ and differential gene expression analysis was performed with DESeq2 ^19^, using a negative binomial distribution algorithm to recognize genes with a p-value ≤ 0.05 and fold change ≥ 1.5. The volcano plot was generated with RStudio (version 4.4.1).

### Transgene Construction

The plasmid pSBtetTA-YP_CFP was built from pSBtet-GP (Addgene #60495)^16^. An EcoRI site was generated (5’-gag to gaa) at eGFP E223 via site directed mutagenesis (NEB #E0554) with primers 5’-tcctgctggaAttcgtgaccg and 5’-ccatgtgatcgcgcttctcg. A Eco81I/ EcoRI-flanked Yellow Fluorescent Protein (YFP) fragment (M plus V1..L221) was generated from Venus BBa_J176006 with primers 5’-tctgcacctgaggccaccatggtgagcaagggcgagg and 5’-ggtcacgaattccagcaggaccatgtgatcg via high fidelity PCR followed by spin-column purification (NEB #E0555, Sigma #NA1020). The YFP fragment and mutated pSBtet-GP plasmid were double-digested with Eco81I/ EcoRI (Thermo #FD0374, #FD0274), gel-purified (NEB #T1020), and ligated (NEB #M2200 with 10x ligase buffer NEB #B0202S) to build pSBtetTA-YP_luc. An SfiI-flanked CFP+NLS fragment (M1..R231 plus nuclear localization signal PKKKRKV) was generated from AmCyan-NLS in BBa_S04698 with primers 5’-tgaaggcctctgaggccaccatggcgctgtccaacaag and 5’-gcttggcctgacaggccttataccttgcgctttttcttggg via high fidelity PCR followed by spin-column purification. SfiI-digested CFP+NLS was ligated into SfiI-linearized pSBtetTA-YP to replace luciferase. Transformations were done with NEB Turbo cells (NEB #C2984) in 100 µg/mL ampicillin selection media without heat shock or recovery.

### Transgenic Cell Lines

DNA lipoplexes were formed with 1.3 μg pSBtetTA-YP_CFP plasmid, 0.1 μg helper SB100X plasmid ^16^ (9:1 molar ratio of pSBtetTA-YP_CFP to SB100X), 5 μL Lipofectamine LTX, and 2.5 uL PLUS Reagent in Opti-MEM in a final volume of 500 μL. Lipoplexes were added dropwise to 2×10^5^ BT-549 or HEK293 cells in 2 mL growth medium (without pen-strep) in 6-well plates, incubated for 1 to 2 days, and inspected for YFP expression. Cells were harvested and expanded to 90% confluency in selection media (0.5 μg/mL puromycin) in 10 cm plates. Sub-samples (5×10^4^ cells) were induced with 0.5 μg/mL doxycycline in a separate 6-well plate to confirm CFP expression. The polyclonal transgenic cells were serial diluted 2-fold in a 6-well plate, grown until single colonies were visible. Colonies were incubated with 0.25% Trypsin buffer at room temperature for 5 minutes, transferred with a 200 μL micropipette into a 24-well plate with standard growth medium, and expanded to ∼90% confluency prior to further outgrowth, LN2 storage, and downstream assays.

### Fluorescent Imaging with Microscopy

Live or fixed and stained cells were imaged in tissue culture plates on an EVOS M5000 inverted microscope (Thermo #AMF5000) at 40x, 100x or 200x magnification. Channel settings were as follows: RGB - no light cube, bright field; YFP - ex. 500/24, em. 542/27 (Thermo #AMEP4954); CFP - ex. 445/45, em. 510/42 (Thermo #AMEP4953). Image processing was done with FIJI/ ImageJ (version macOS x86_64).

### Flow Cytometry

A minimum of 5×10^4^ cells were prepared and analyzed per run on a CytoFLEX Flow Cytometer as follows. Cells were harvested, washed with 1X DPBS, pelleted, and resuspended in FACS Buffer (10% FBS and 5 mM EDTA in 1X DPBS) on ice. Transgenic BT549 cells treated with 1 μg/mL or 0 μg/mL of doxycycline were used as positive controls for CFP (K0525 V 525/40 Laser; ex. 468, em. 498) or YFP (FITC B 525/40 Laser ex. 498, em. 517) channels, respectively. Parental (nontransgenic) cells were used as non-fluorescent controls. Viability was detected using Zombie NIR (BioLegend #423105) staining on live and heat-killed parental BT549 cells. Data was visualized in real time using CytExpert software, and statistically analyzed using FlowJo software (version 10.1).

## RESULTS

### Tet-TA regulated transgene expression is perturbed in ACM-treated, lipogenic TNBC cells but not in HEK293 cells

To model biochemical signaling from adipocytes to breast cancer epithelial cells, we cultured differentiated mouse OP-9 cells, collected the adipocyte conditioned medium (ACM) after 3 days, and treated breast epithelial BT-549 cells with 50% ACM (**Figure 1A**). Three days after treatment, we observed a striking accumulation of lipid droplets in the cytoplasm of most cells, whereas lipid droplets were barely visible or absent in cells grown in unconditioned medium (UCM) (**Figure 1B**). These results are consistent with other reports where adipocyte-secreted factors induce lipid accumulation in breast cancer cells ^20,21^ and are thought to serve a cell-protective, pro-oncogenic role. Therefore, adipocyte secreted factors in ACM collected from mouse cells can induce a lipogenic state in human TNBC cells *in vitro*.

**Figure 1.**
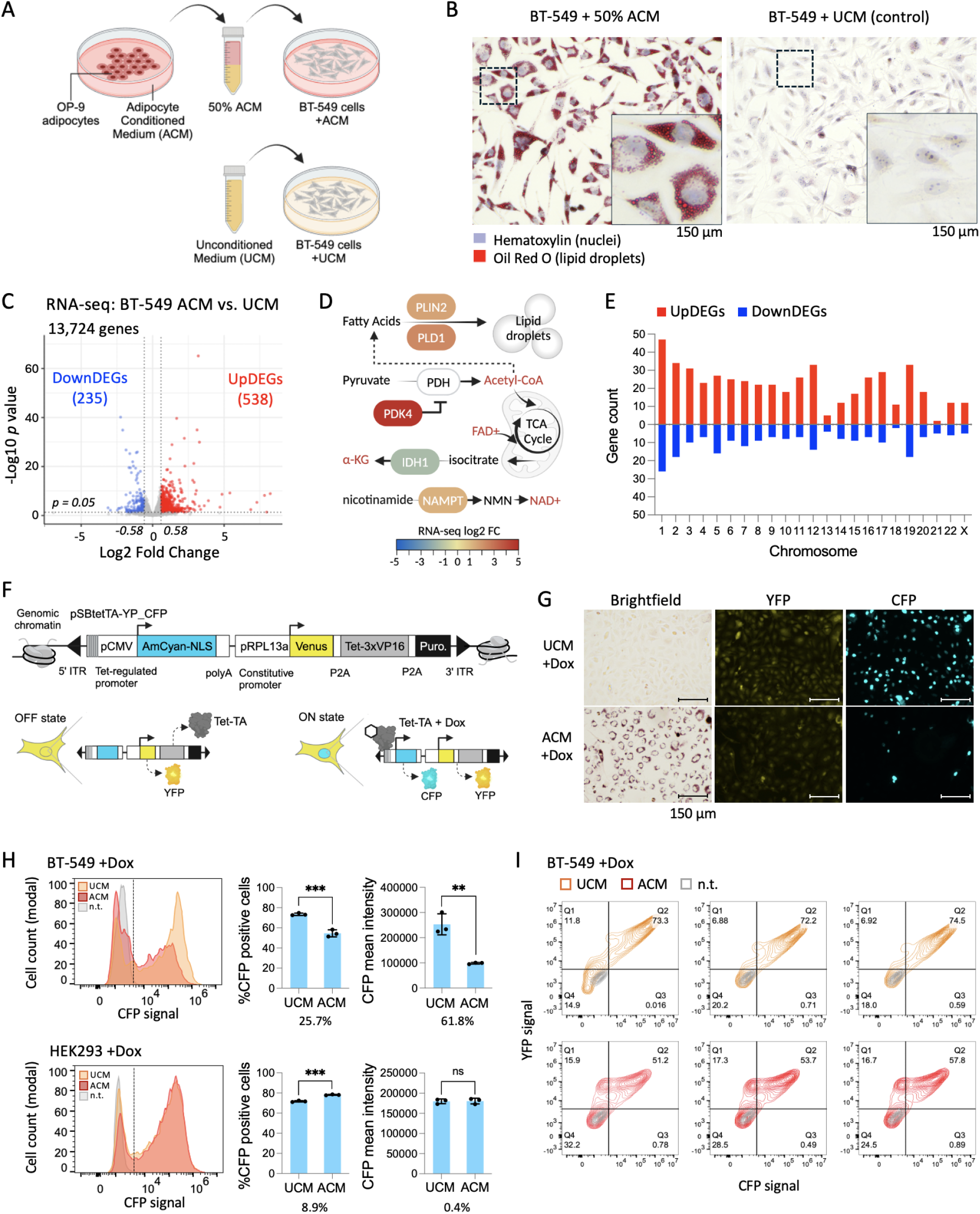
Adipocyte conditioned medium induces a lipogenic state accompanied by the repression of endogenous genes and a transgene in BT-549 breast cancer cells. A. OP-9 mouse stromal cells are differentiated *in vitro* and allowed to secrete factors including cytokines and free fatty acids into the culture medium. The adipocyte conditioned medium (ACM) is collected and added to cultured TNBC cells.B. Oil Red O staining of BT-549 cells grown in unconditioned medium (UCM, control) or 50% ACM for three days. C. RNA-seq expression profiling of ACM-treated cells compared with UCM-treated cells. D. Metabolic functions of enzymes expressed by select differentially expressed genes (DEGs). Epigenetic substrates and cofactors are shown in red text. E. Genomic distribution of up- and down-regulated DEGs (UpDEGs, DownDEGs). F. The chromosomally integrated transgene pSBtetTA-YP_CFP contains a doxycycline-inducible CMV promoter that drives expression of a AmCyan fluorescent protein tagged with a nuclear localization sequence (CFP-NLS), and a constitutive RPL13a promoter that drives expression of Venus yellow fluorescent protein (YFP), the Tet-3xVP16 activator, and puromycin resistance. G. Fluorescence imaging of CFP-NLS expression in UCM (control) and 75% ACM-treated BT-549 cells. H. Flow cytometry analysis of CFP-NLS expression in UCM or 75% ACM-treated BT-549 TNBC cells and HEK293 kidney cells. Histograms show one representative sample from each condition. Bar charts show means of three replicate wells per condition, percent differences for mean UCM versus mean ACM, standard deviation (black bars), and unpaired t-test values: *p* ≤ 0.05*, 0.01**, 0.001***, or not significant (ns). I. Contour plots of YFP and CFP flow cytometry signals from three replicates per condition. Illustrations (A, D, F) were generated in BioRender.

Acetyl-CoA is a major cellular metabolite that is used to synthesize fatty acids and triacylglycerols for energy storage ^22^, and is also required for histone acetyltransferase activity in the nucleus, which generally supports active gene transcription. We investigated whether increased lipogenesis is sufficient to impact gene regulation. We hypothesized that during lipogenesis, a lack of available acetyl-CoA would lead to the loss of expression at many endogenous genes. Comparative RNA-seq transcription profiling of BT-549 cells cultured in 50% ACM versus UCM identified 235 significantly repressed genes (DownDEGs), as well as 538 activated genes (UpDEGs) (fold change threshold 1.5, *p* ≤ 0.05, **Figure 1C**). Consistent with the lipogenic phenotype, key lipid droplet biosynthesis genes became upregulated: *PLD1* (phospholipase D1) ^23^ and *PLIN2* (perilipin-2) ^24^ (**Figure 1D**). Other metabolic enzymes encoded by DEGs included PDK4 (pyruvate dehydrogenase kinase 4) which inhibits PDH ^25,26^, IDH1 (isocitrate dehydrogenase 1) which converts isocitrate into α-KG ^27^, and NAMPT (nicotinamide phosphoribosyltransferase) which is the rate-limiting component in the NAD biosynthesis pathway ^13^. High PDK4 and low IDH1 might lead to decreased levels of acetyl-CoA and α-KG needed for the formation of transcriptionally active chromatin ^28^. High NAMPT might increase NAD+ levels to support transcriptionally repressive chromatin through Sirtuin-mediated histone deacetylation. DownDEGs appeared at every chromosome (**Figure 1E**), suggesting that epigenetic repression occurred at loci distributed throughout the genome.

To investigate the consequence of this broad gene regulation on transgene behavior, we generated stable transgenic cell lines using a customizable sleeping beauty transposon-based reporter that allows semi-random insertion via homology with repetitive sequences across the genome^16^. The constitutive expression of Venus yellow fluorescent protein (YFP), a Tet-3xVP16 transcriptional activator (Tet-TA), and puromycin are driven by the RPL13a promoter. Inducible expression of AmCyan fluorescent protein (CFP) tagged with a nuclear localization sequence (CFP-NLS) is regulated by doxycycline-bound Tet-TA, which binds to sequences upstream of a CMV promoter (**Figure 1F**). Non-lipogenic cells grown in UCM showed dim to very bright nuclear CFP after treatment with 1.0 μg/mL doxycycline for two days. Cells co-treated with 75% ACM and doxycycline for two days showed reduced activation of CFP-NLS (**Supplemental Figure S1**), and repression was even more pronounced in cells treated with 0.5 μg/mL doxycycline (**Figure 1G**). Visual inspection showed that low CFP-NLS signal generally corresponds with high lipid droplet content in single cells (**Figure 1G**).

Quantitative analyses were carried out with flow cytometry to further investigate changes in transgene expression in response to ACM. ACM treatment reduced CFP-positive cells by 25.7%, and the mean fluorescence intensity of CFP-positive cells was reduced by 61.8% compared to the UCM samples (**Figure 1H)**. To determine if this phenomenon is cell type specific, we generated HEK293 cells that carried the same construct and subjected these cells to the same experiments. CFP-NLS expression showed either modest changes (less than 10%) or no significant changes in one transgenic cell line (**Figure 1H**) and in additional clonal HEK293 isolates (**Supplemental Figure S2**). We did not observe substantial changes in cell morphology or lipid droplet content in ACM-treated HEK293 cells, suggesting that ACM-induced lipogenesis and epigenetic repression are cell-type dependent.

It is possible that loss of the Tet-TA regulator via epigenetic silencing of the *pRPL13a* promoter contributes to the loss of CFP. To investigate this idea we compared the expression of YFP, which is also regulated by *pRPL13a* (**Fig. 1F**), in the UCM +Dox and ACM +Dox samples. 15.9% - 17.3% of the cells were YFP-positive and CFP-NLS-negative (**Fig. 1I**), suggesting that in some cells the inducible *pCMV* promoter that drives CFP-NLS is distinctly, more strongly repressed by the lipogenic epigenetic state.

### Activation of *pCMV* prior to ACM treatment protects the transgene from full repression by the lipogenic epigenetic state

Our previous experiments showed different effects of the lipogenic state on each of the two fluorescence-expressing genes in pSBtetTA. This raises the possibility that strong active transcription, i.e. from *pRPL13a*, is less affected by the lipogenic state and that transcriptional initiation, i.e. via Tet-TA binding to *pCMV*, may be more sensitive to repression. To investigate the sensitivity of transcription initiation to the lipogenic state, we treated cells with 75% ACM for one day, and then activated CFP expression by adding doxycycline to the samples (**Figure 2A**, Treatment 1). We used the data from our previous ACM and doxycycline co-treatment experiments for comparison (Treatment 2). Additionally, to determine if active transcription protects CFP from repression, we treated cells with doxycycline to activate expression for two days before treating the cells with 75% ACM (Treatment 3). We waited at least 48 hours after ACM treatments before imaging the cells, which we expected to be sufficient for fluorescent protein degradation after transcriptional silencing ^29^.

**Figure 2.**
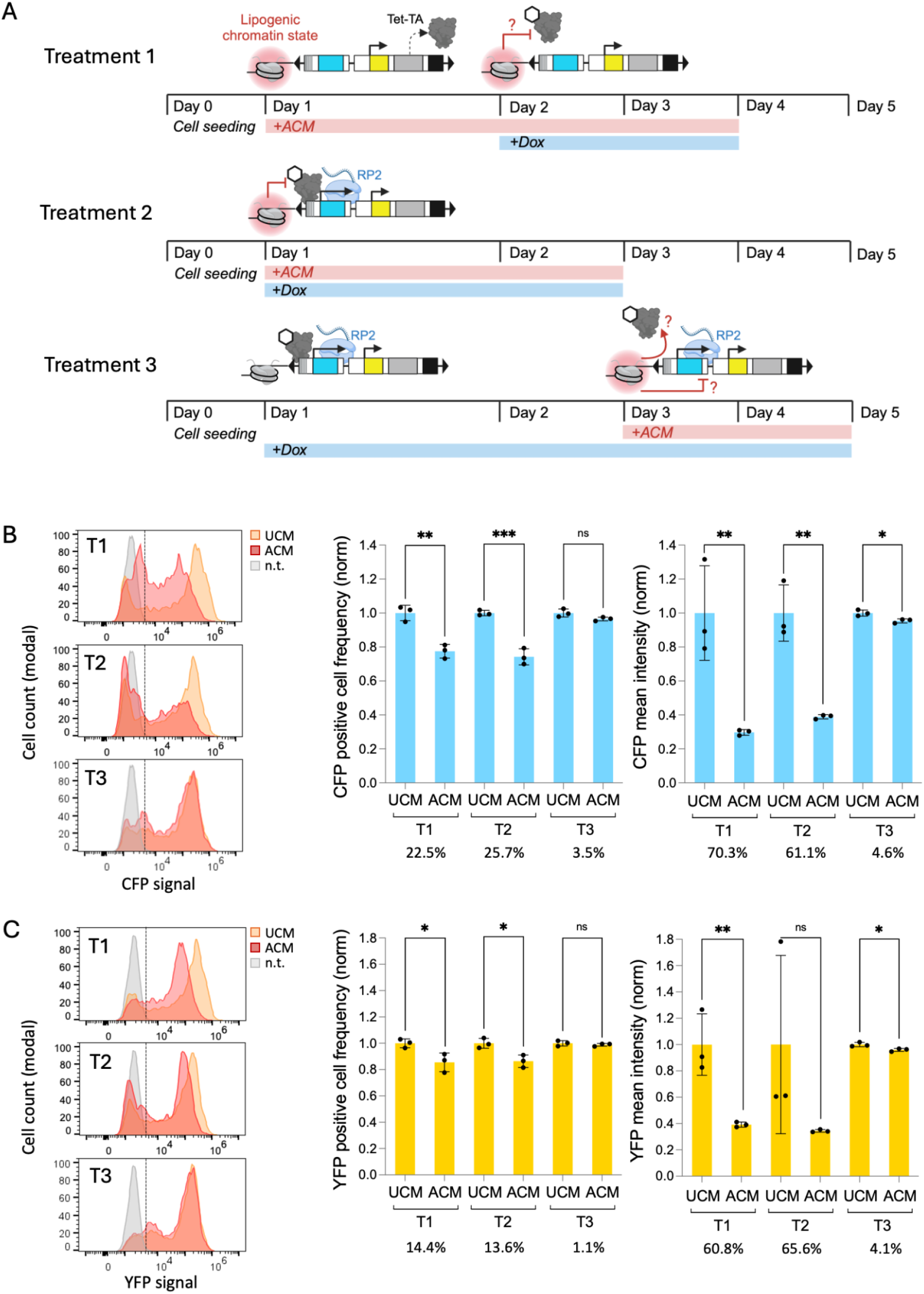
Comparison of lipogenic epigenetic repression of pSBtetTA-YP_CFP before, during, or after *pCMV* activation. A. Experimental design. Controls were included in each treatment (T1, T2, and T3), where the media was replaced with fresh UCM instead of ACM. B. Flow cytometry quantification of CFP expression. N.t. = non-transgenic BT-549 cells. C. Quantification of YFP expression in the samples from panel B. Bar charts show values normalized by the mean of three UCM replicates within each group (T1, T2, or T3), percent differences for mean UCM versus mean ACM, standard deviation (black bars), and unpaired t-test values: *p* ≤ 0.05*, 0.01**, 0.001***, or not significant (ns).

Pre-initiation of the lipogenic state (**Figure 2B**, Treatment 1) in BT549 cells resulted in a 22.5% decrease in the percentage of CFP-positive cells and a 70.3% decrease in the mean intensity of CFP. The dual initiation of the lipogenic state with *pCMV* activation (**Figure 2B**, Treatment 2) resulted in a 25.7% decrease in the percentage of CFP-positive cells and a 61.1% decrease in the mean intensity of CFP expression. Inducing the lipogenic state before *Pcmv* activation (Treatment 1) resulted in a 8.9% stronger decrease in CFP mean intensity compared to Treatment 2. This suggests that repressive epigenetic mechanisms of the lipogenic state prevent transcriptional initiation, possibly by perturbing Tet-TA binding to TetO sequences at *pCMV*. Conversely, the pre-initiation of *pCMV* activation (**Figure 2B**, Treatment 3) resulted in a small, insignificant decrease in the percentage of CFP-positive cells and only a 4.6% decrease in the mean intensity of CFP expression. For Treatment 3, *pRPL13a*-driven expression of YFP was also modestly affected. These results suggest that active transcription protects the pSBtetTA transgene from lipogenic epigenetic silencing.

Next, we compared changes in *pRPL13a*-driven YFP expression to changes in Tet-TA/*pCMV* regulated CFP expression. Across all treatment regimes, YFP-positive cell frequency was less affected, showing smaller percent differences than the CFP-positive cell frequencies (**Figure 2C**). This is consistent with our observation that some ACM-treated cells that were CFP-negative were also YFP-positive (**Figure 1I**). In contrast, mean YFP signal intensity was reduced similarly to CFP with pre-initiation of the lipogenic state (Treatment 1) and dual-initiation of the lipogenic state and *pCMV* activation (Treatment 2). It is worth noting that, removing the high YFP outlier in the Treatment 2 group would show that YFP mean intensity is less affected than for Treatment 1. The decrease in mean YFP intensity suggests epigenetic suppression of *pRPL13a* activity, which drives Tet-TA expression. Therefore potentially lower concentrations of the Tet-TA regulator might contribute to the loss of CFP expression. Although YFP levels were reduced, most cells maintained detectable expression, supporting the idea that active transcription can protect the transgene from full silencing by the lipogenic epigenetic state.

## DISCUSSION

Our results demonstrate how the shift in a metabolic state that affects levels of metabolites involved in chromatin remodeling can influence transgene expression in engineered human cells. Activation of lipogenesis-associated genes is driven by mTOR-activated master transcriptional regulators such as SREBP. Genes that lack SREBP target sequences and other mTOR-activated regulatory elements might be prone to epigenetic repression through the impairment of chromatin remodeling enzymes that rely on metabolites that are consumed or decreased during lipogenesis. Lipogenesis consumes acetyl-CoA as a building block for fatty acid synthesis, thus less acetyl-CoA is available for the TCA cycle. TCA cycle slow-down reduces the production of α-KG. Transgene repression might be the consequence of impaired chromatin remodelers that support transcriptionally-active chromatin, including acetyl-CoA-dependent histone acetyltransferases (HATs), and α-KG-dependent enzymes that remove methyl groups from DNA and histone H3 lysine 9 and 27 ^28^. Certain chromatin modifiers that support repressive chromatin, such as enzymes that demethylate histone lysine 4, also require α-KG, suggesting that chromatin silencing might also be disrupted. However, the formation of repressive chromatin might be reinforced through several mechanisms. For instance, increased expression of NAMPT might increase NAD+ which is required for class III histone deacetylase (Sirtuin) activity. Furthermore, NAD+-independent histone deacetylase 5 (HDAC5) ^30,31^ became upregulated in ACM-treated BT-549 cells (FC = 1.9, *p* < 0.001). Finally, TCA cycle slow-down might slow the consumption of FAD+, possibly making this cofactor available for enzymes that add methyl marks to DNA and histone lysine 9 and 27.

Our results also suggest that the sensitivity of the transgene to the lipogenic epigenetic state might depend upon the initial state of the promoter. We showed that when lipogenesis is stimulated prior to activation of *pCMV*, the mean expression level was decreased. In contrast, pre-activation of the promoter appeared to protect the same promoter from full repression. These results suggest a mechanism where the initial activation of the promoter establishes a favorable chromatin environment, potentially marked by histone acetylation and persistent open chromatin that resists the effects of reduced acetyl-CoA and α-KG.

One key limitation of our study is the reliance on *pCMV* and *pRPL13a* as model promoters. While *pCMV* is a widely used promoter for cell engineering, other promoters may respond differently to metabolic states or may be more susceptible to repression due to sequence-specific interactions with other transcription factors. Future work could explore the behavior of tissue-specific or inducible promoters under varying metabolic conditions to deepen insights into transgene behavior across different cellular contexts. Furthermore, our study did not explicitly analyze chromatin accessibility through techniques like ATAC-seq or ChIP-seq, which could provide direct evidence of epigenetic changes associated with transgene repression. Incorporating such analyses could help identify specific histone modifications and DNA methylation patterns responsible for the observed transcriptional outcomes.

These results have implications for understanding transgene behavior in tissue microenvironments (*in vivo*) undergoing metabolic changes, and for optimizing *in vitro* cell culture to avoid transgene silencing. Furthermore, the fluorescent reporter system used here is a potentially useful tool for tracking epigenetic states at the single-cell level in biophysically relevant models such as mouse cancer xenografts and adipocyte-epithelial cell co-cultures. Our observations also have implications for transgene-based cell engineering beyond cancer models. The shift between glucose-dependent and lipid-dependent energy production, glycolysis and lipolysis, in response to nutrient availability and cell signaling is an essential process in industrially important eukaryotic cells including yeast, chinese hamster ovary (CHO) cells, and human stem cells undergoing differentiation and maturation. Therefore scientists might consider monitoring the expression levels of key enzymes involved in lipogenesis when transgene expression becomes unstable in engineered cells.

## CONCLUSION

This work demonstrates that in addition to previously reported cellular factors, transgene repression in cultured human breast cancer cells can occur as a consequence of a lipogenic state induced by adipocyte-secreted factors. Using a customizable sleeping beauty (pSB) based transgenic system, we observed lipogenesis-associated repression of a Tet-TA-targeted CMV core promoter of a cyan fluorescent reporter gene. These results link metabolic changes that affect cofactors and substrates of chromatin remodeling enzymes to transgene expression. This insight will support strategies to optimize transgene expression in *in vitro* and *in vivo* models.

## ACKNOWLEDGEMENTS

This study was supported by grants from the University of Michigan/ Genentech Research Awards Program (GRAP), PureTech, and Developmental Funds from the Winship Cancer Institute of Emory University. This study was supported in part by the Emory Integrated Core Facility (EICF) Emory plus Pediatrics/ Winship Flow Cytometry Core (ECFCC) award number P30CA138292 (to Emory University). The content is solely the responsibility of the authors and does not necessarily represent the official views of the National Institutes of Health.

