## Supplemental Information for "Tet Transgene Activation is Disrupted in Lipogenic Triple Negative Breast Cancer Cells"

Supplemental Figures

- Figure S1. Comparison of the CFP-off state with CFP-on states in UCM and ACM-treated BT-549 cells.
- Figure S2. Transgene expression in HEK293 cells treated with UCM or ACM.

Supplemental Methods

- Preparation of Insulin Oleate Medium (IOM) for OP9 differentiation

**SUPPLEMENTAL FIGURES**

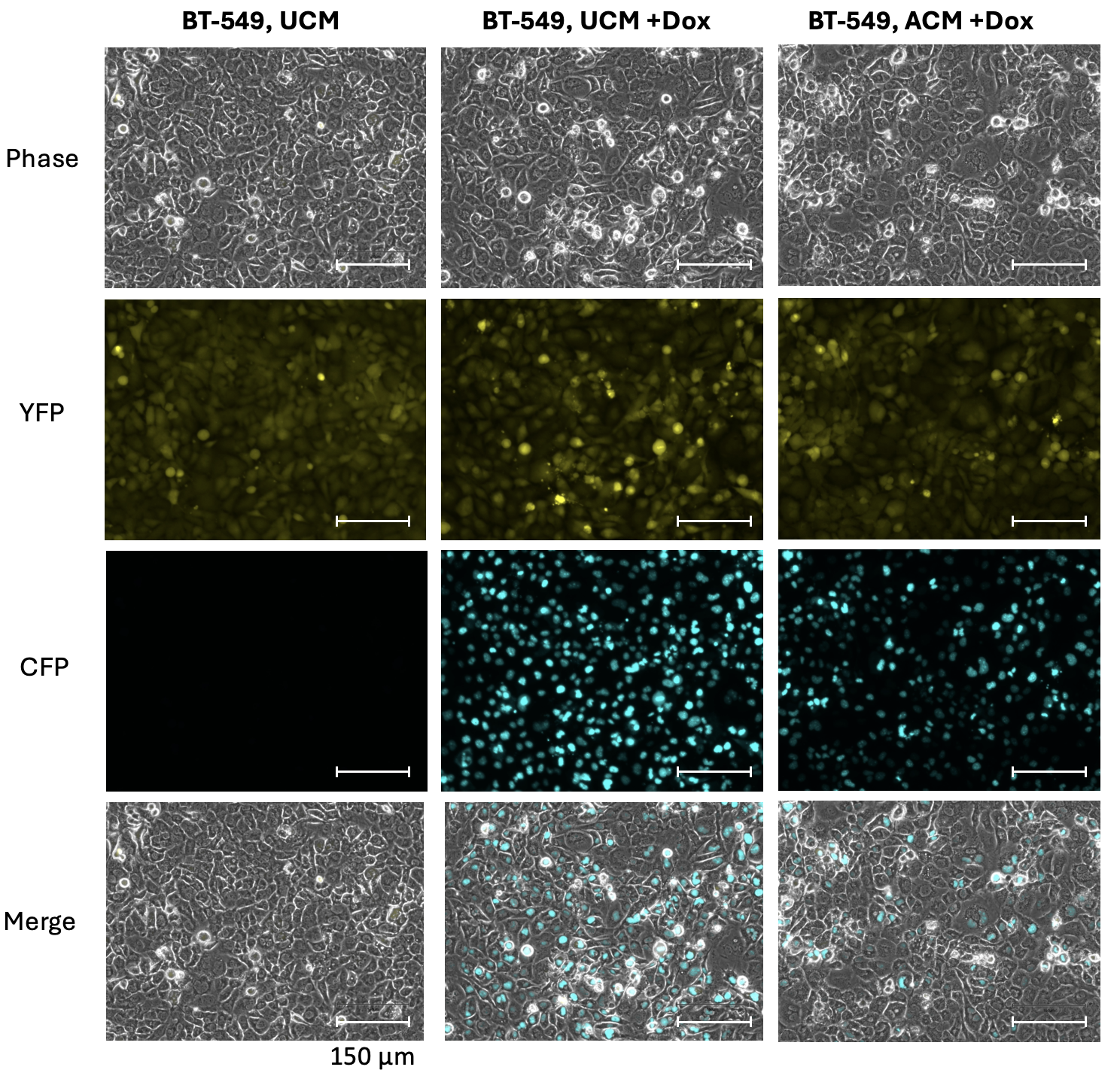

**Figure S1.** **Comparison of the CFP-off state with CFP-on states in UCM and ACM-treated BT-549 cells.** Cells were seeded in 6-well plates, grown in unconditioned medium (UCM), UCM plus 1.0 μg/mL doxycycline (Dox), or 50% adipocyte conditioned medium (ACM) plus 1.0 μg/mL Dox and imaged after two days.

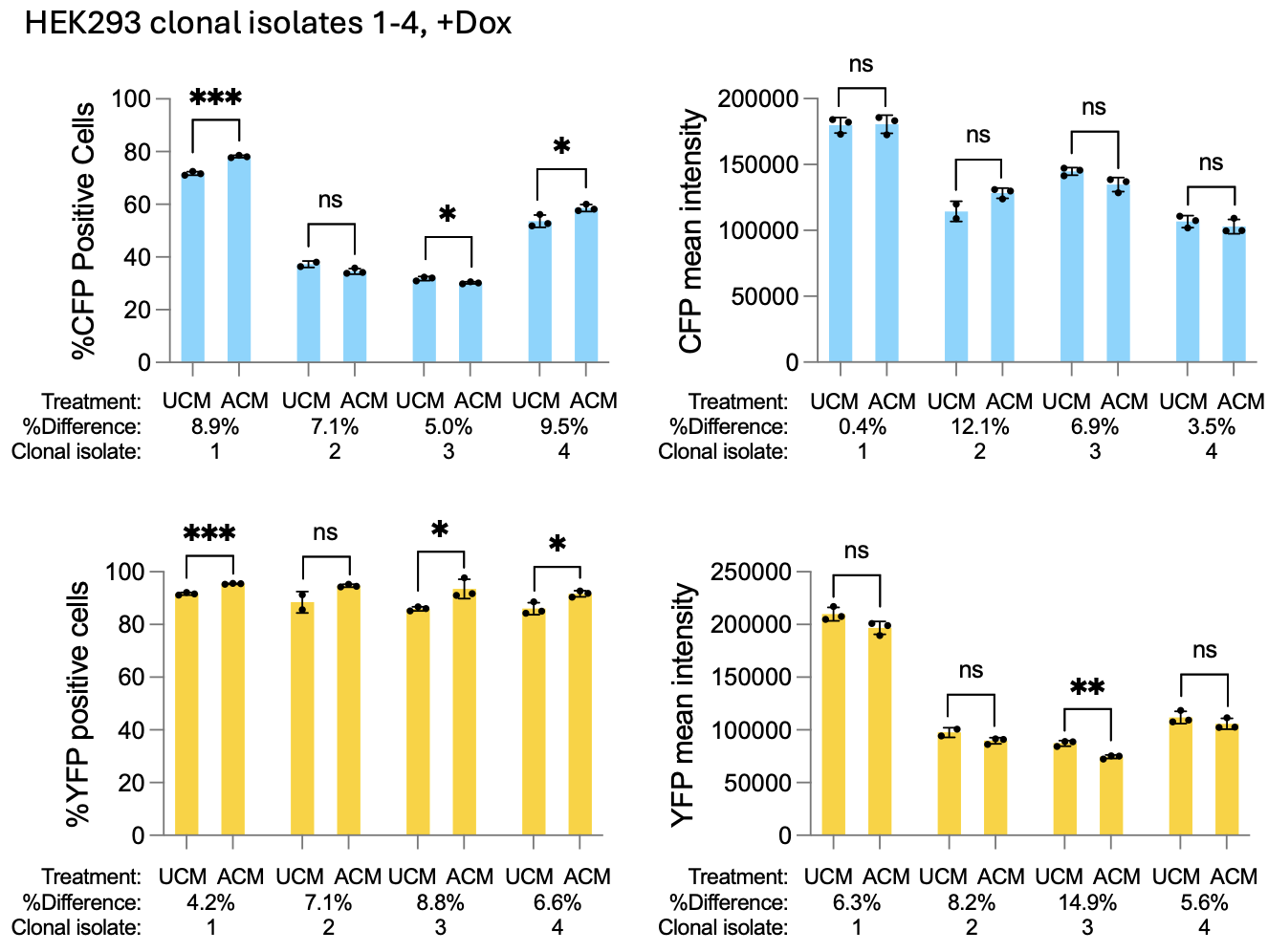

**Figure S2. Transgene expression in HEK293 cells treated with UCM or ACM.** Bar charts show means of three replicate wells per condition, percent differences for mean UCM versus mean ACM, standard deviation (black bars), and unpaired t-test values: *p* ≤ 0.05*, 0.01**, 0.001***, or not significant (ns).

**SUPPLEMENTAL METHODS**

**Preparation of Insulin Oleate Medium (IOM) for OP9 differentiation**

A 100 mM stock solution of Sodium Oleate was prepared with 100% methanol and a stock solution of 30% fatty acid free BSA was prepared with 1X PBS. Both reagents were passed through a 0.22 μm filter before use. The Insulin Oleate Media (IOM) culture media was prepared with 1.52 mM Sodium Oleate (Sigma #O7501) and 1.8% fatty acid free BSA (Sigma t#A6003) and incubated at 37°C for 2 hours. After incubation, the IOM media was supplemented with 0.2% FBS, 175 nM insulin (Sigma #I6634), and 1% Penicillin-Streptomycin (Thermo Fisher Scientific #15140122). The final volume of IOM was made up of alpha-MEM media (Gibco #12561056).
